# Experimental evidence for the benefits of higher X-ray energies for macromolecular crystallography

**DOI:** 10.1101/2021.01.21.427633

**Authors:** S. L. S. Storm, D. Axford, R. L. Owen

## Abstract

X-ray induced radiation damage is a limiting factor for the macromolecular crystallographer and data must often be merged from many crystals to yield complete datasets for structure solution of challenging samples. Increasing the X-ray energy beyond the typical 10-15 keV range promises to provide an extension of crystal lifetime via an increase in diffraction efficiency. To date however hardware limitations have negated any possible gains. Through the first use of a Cadmium Telluride Eiger2 detector and a beamline optimised for high energy data collection, we show that at higher energies fewer crystals will be required to obtain complete data, as the diffracted intensity per unit dose increases by a factor of more than 3 between 12.4 and 25 keV. Additionally, those higher energy data provide more information, evidenced by an increase in high-resolution limit of up to 0.3 Å, pointing to a high energy future for synchrotron-based macromolecular crystallography.

## Introduction

Synchrotron-based macromolecular crystallography (MX) is the method of choice for determining the atomic structure of proteins and viruses, providing almost 90% of Protein Data Bank depositions over the last 5 years^1^. More than 95% of these synchrotron-derived depositions were collected using X-rays with energies in the range 10-15 keV (1.240 – 0.827 Å) reflecting the optimisation of sources, beamlines, and detectors within this narrow region and, on the sample side, the success of seleno-methionine incorporation for experimental phasing at 12.67 keV ^2^. The continual development of synchrotron beamlines and sources has resulted in the realization of smaller beam sizes and increased flux densities at the sample position^3^. While these brighter beams enable structure solution from ever-smaller and more challenging crystals it is at the expense of the one-crystal one-structure approach as X-ray induced damage precludes the collection of a complete dataset from a single crystal^4^. In such cases, formation of a complete dataset is achieved using a multi-crystal methodology, distributing the total dose required for structure solution over many crystals^5,6^.

Robust approaches for both collecting and processing multi-crystal data have been developed^5,7–11^ with the logical endpoint being serial synchrotron crystallography where a single diffraction image is collected from each crystal^12^. Rather than collecting from ever more crystals however a primary aim of a multi-crystal experiment should be to optimize the last experimental step, maximising the volume of data that can be collected from each crystal, reducing sample consumption and simplifying data collection and subsequent analysis.

Increasing the energy of the incident X-rays as a solution to the multi-crystal challenge is attractive as no change to, or treatment of, the crystals used is required to achieve the change and the approach is universally applicable as it exploits the differing energy dependence of how X-rays interact with matter via elastic scattering, inelastic scattering or the photoelectric effect. The resulting benefits of higher energies are two-fold. Firstly, as the X-ray energy increases the number of elastically scattered photons per unit absorbed dose, or Diffraction Efficiency (DE), increases which is reflected experimentally in higher diffraction intensities for a given dose (improved I/D ratio)^13–15^ meaning crystal exposures can be reduced. Secondly, at higher energies photoelectron escape means energy deposited by X-rays can leave the crystal, depending on its volume, further reducing the absorbed dose^16^. A key result from many years of work on radiation damage to cryo-cooled crystals is that X-ray induced damage is proportional to the absorbed dose^17^. Both of the effects introduced above predict higher diffracted intensities per unit absorbed dose at higher energies thus the dose can be reduced to obtain the same diffraction intensities. Consequently, the use of higher X-ray energies implies that more useful diffraction data can be collected from each crystal.

The energy dependence of DE was first noted by Arndt (1984), who showed that for crystalline proteins the probability of photoelectric absorption decreases more rapidly with increasing photon energy than the probability of elastic scattering. The intensity of Bragg spots can be predicted using Darwin’s equation^18^, which takes X-ray beam and crystal parameters into account. A closer inspection of this also reveals a resolution dependence^19^ with gains in DE at higher X-ray energies enhanced for higher resolution reflections.

In addition to the decrease in photoelectric absorption at high energies, another effect decreasing the deposited dose at higher energies is photoelectron escape. Nave & Hill (2005) simulated the track of photoelectrons in micro-crystals and concluded that a significant proportion of photoelectrons could leave the crystal before causing damage ^16^. The inclusion of photoelectron escape and Compton scattering into calculation of the diffraction efficiency shows a theoretical five-fold gain in DE for 5 μm crystals when the energy of incident X-rays is increased from 7 to 30 keV^20^.

To date attempts to demonstrate the benefits of high energy data collection on protein crystals have been hampered by an absence of suitable detectors. Most experiments performed at high energies have been performed with CCD detectors^14,21,22^ partially combined with X-ray image intensifiers ^23^, image plates ^21,24–26^ or even with point detectors^27^. All studies performed with two-dimensional detectors report that the detective quantum efficiency is rather low and not well characterized for high energies. Currently, the most widely applied detector technology for recording diffraction data at synchrotrons utileses a hybrid photon counting (HPC) approach. HPC detectors feature a sensor bonded to an electronic counter and X-ray photons absorbed by the sensor material are recorded as a ‘count’ ^28^. Typically the sensor material is silicon, a widely available, easily processible material. As the atomic number of silicon is low, the sensor rapidly becomes transparent as the X-ray energy is increased beyond 15 keV however. This decrease in absorption means the quantum efficiency (QE) of silicon-based detectors falls rapidly as a function of energy: to less than 20% at 25 keV for a 450 μm thick Si sensor ^29^. Recently, detectors using cadmium telluride as a sensor material have been developed; the use of CdTe results in a detector QE of more than 90% below the cadmium absorption edge (26.7 keV) and nearly 80% up to energies of 80 keV^30^. Simulations with RADDOSE-3D taking into account the quantum efficiency for a detector with a 750 μm thick CdTe sensor and assuming a top-hat beam profile predict an optimal data collection energy of 26 keV ^31^.Initial experiments with a Pilatus3 detector equipped with a CdTe sensor were recently successfully used in MX to experimentally prove the benefits of photoelectron escape in micro-crystals^32^, to collect data at 35 keV to ultra-high resolution^33^ and to investigate specific radiation damage at high energies^34^.

Here we present the first use of a CdTe Eiger2 detector for routine high energy MX. The detector provides a nearly constant quantum efficiency across the investigated energy range. We experimentally demonstrate increased dose efficiency at higher energies observing an almost four-fold increase between 12.4 and 25 keV. No gain is observed when an identical experimental approach is employed using a silicon detector. We further observe an increase in the resolution of data obtained for a given absorbed dose at higher energies. The combination of these effects will allow fewer crystals to be used for structure determination and our results point to a high energy future for synchrotron-based MX.

## Results and Discussion

In order to quantify the energy dependence of the diffraction efficiency of protein crystals, 29 low-dose data series were collected from 11 thermolysin crystals. Each data series consisted of four 100° wedges collected from the same position on a crystal with each wedge recorded using a different X-ray energy (12.4 keV, 17.5 keV, 22.3 keV or 25 keV). 22 data series were recorded using an Eiger2 9M (750 μm thick CdTe sensor) and 7 with a Pilatus3 6M (450 μm thick Si sensor) detector. In order to minimize the contribution of radiation damage to observed trends, the total absorbed dose was kept low: to less than 800 Gy per sweep *i.e.* less than 2 % of the 43 MGy required to half the diffracting power of cryo-cooled crystals^35^ and the order of energies varied between each series (table S1).

Figure 1 shows the agreement between the observed diffracted intensity divided by the beam intensity (I/I_beam_) as a function of energy for the crystals used in this study and I/I_beam_ predicted by Darwin’s equation which is presented in a simple form by Giocovazzo^36^

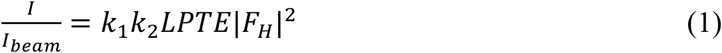

**Figure 1.**
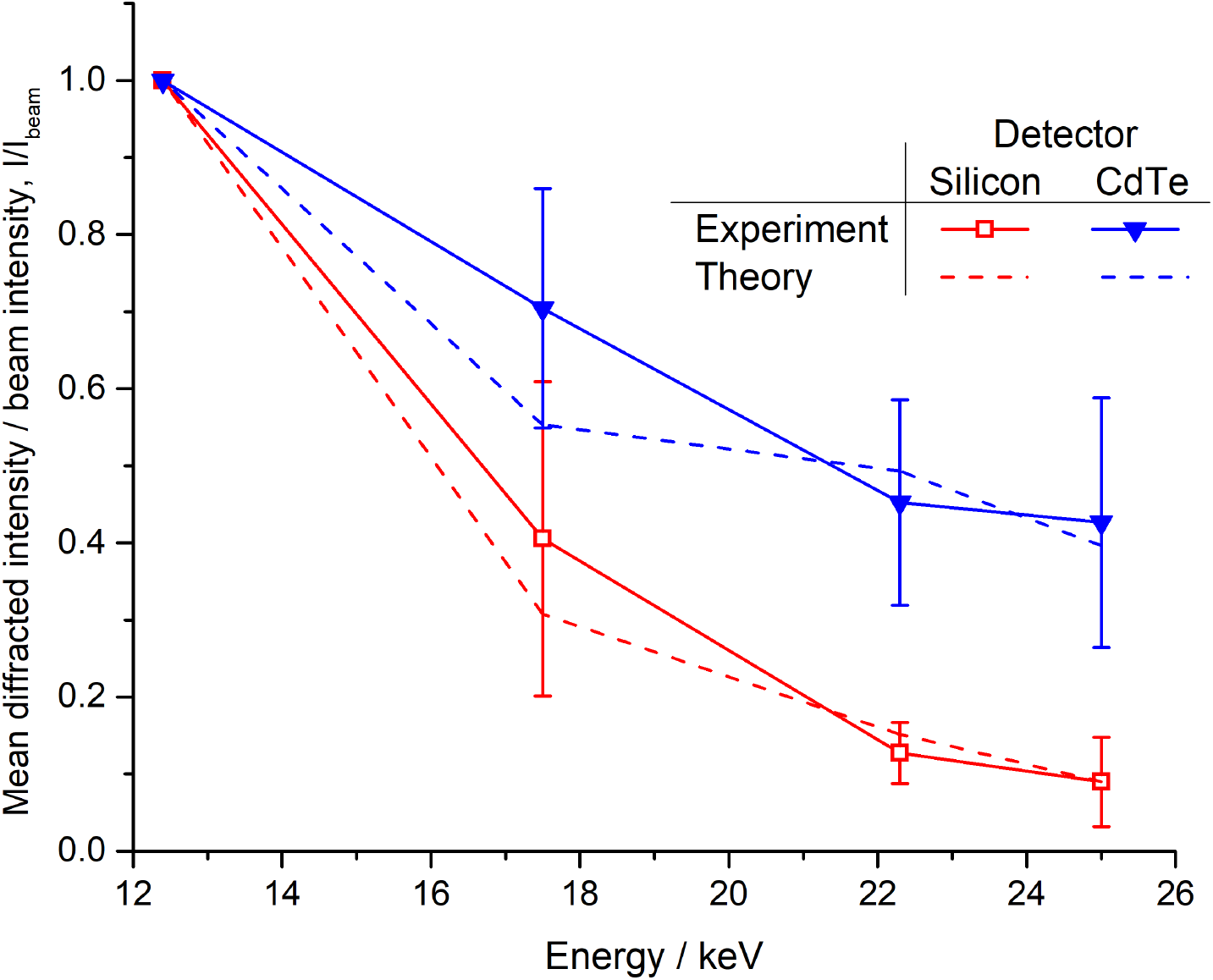
Ratio of diffracted intensity to beam intensity (I/I_beam_) as a function of energy for data recorded using both the CdTe Eiger and Si Pilatus detectors. Experimental data are the mean of 22 and 7 data sets for the Eiger and Pilatus detectors respectively. Overlaid is the predicted energy dependence of I/I_beam_ determined from theory using measured fluxes and beamsizes.

Here *k*_1_ = *e*^4^/*m*^2^*c*^4^, takes into account universal constants, and 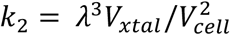 where λ is the X-ray wavelength, *V*_*xtal*_ the illuminated crystal volume and *V*_*cell*_ the volume of the unit cell. *L* is the Lorentz factor, *P* the polarization factor, and *T* and *E* the X-ray transmission and extinction coefficients of the crystal respectively. The analytical expression for the product of 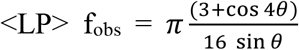 derived by Holton^20^ can be used, and energy independent terms excluded to show

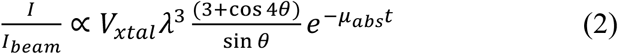

where θ is the Bragg angle, *t* the crystal thickness and *μ*_*abs*_ its X-ray absorption coefficient. When applying this expression to the data collected here we also consider the energy dependent quantum efficiency of the detector used.

The agreement between experiment and theory shown in figure 1 illustrates that the ratio of elastically scattered photons to incident photons varies as expected for both detectors and, importantly, gives confidence in both the accuracy and validity of the quantity mean diffracted intensity as the numerator I in the ratio I/D derived below.

Intensity statistics for two individual data series collected from the same crystal, one recorded using each detector, are shown in figure 2. Both of these data series were collected with the same data collection parameters and the same increasing energy sequence (12.4 keV, 17.5 keV 22.3 keV and finally 25 keV). The mean unscaled intensities recorded over the two data series while aiming at keeping the total diffracted intensity approximately constant are shown in figure 2a. A strong energy dependence is clear for data recorded using the CdTe Eiger detector with higher intensities recorded at higher energies, a trend not observed using the Si Pilatus. The observed signal to noise ratio, <I/σ(I)>, is shown in figure 2b, this illustrates again the advantage conferred by cadmium telluride. CdTe data again show a constant increase as a function of energy whereas the maxima for Si data is at 17.5 keV: at 22.3 and 25 keV when the absorption of silicon is low <I/σ(I)> falls. Concomitant with the decrease in the quantum efficiency of the silicon sensor, the internal consistency of the data also decreases, as can be seen from the increased R_meas_ values for data collected with the Si Pilatus detector at higher energies (figure 2c). We note that other detector properties beyond the sensor material contribute to some of the observed differences: both the smaller pixel size and zero deadtime of the Eiger detector also affect data quality^37^. The 0.95 ms dead time of the Pilatus detector used corresponds to almost 10% of the total exposure time for data collected at energies below 25 keV for example. Our results clearly show however that the CdTe sensor material is essential to exploit the benefits of high energy data collection.

**Figure 2:**
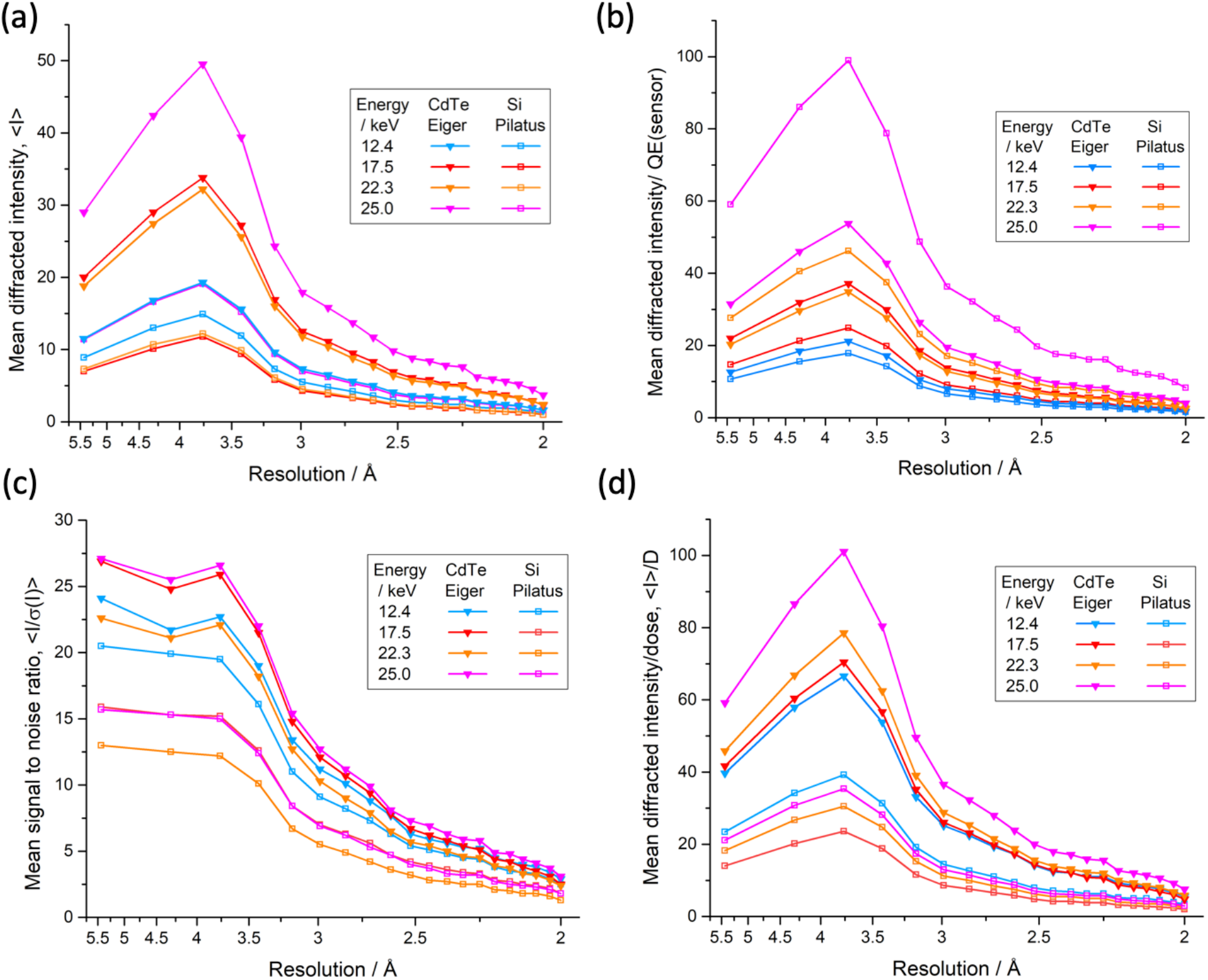
Overview of two data series collected from the same crystal using a CdTe Eiger detector (filled triangles) and a Si Pilatus detector (hollow squares). (a) shows the mean unscaled intensity per Bragg spot <I>, as a function of resolution (b) Mean unscaled signal to noise ratio per Bragg spot <I/σ(I)>. (c) Redundancy independent merging R value R_meas_. (d) Mean diffracted intensity per Bragg spot normalized to the absorbed dose, <I>/D. In all panels data points reflect the high resolution limit of each shell, the lowest resolution data point includes reflections over the range 40 – 5.5 Å.

With doses of less than 540 kGy per data set (table S2) global radiation damage is unlikely to have a significant effect on the observed trends. Indeed 2Fo-Fc maps comparing data sets collected from the same position in the crystal display almost no signs of site-specific radiation damage (figure S1).

Figure 2d shows the diffracted intensity per unit absorbed dose, <I>/D, for the two data series. When the CdTe Eiger detector, optimised for high energy data collection, is used a clear increase in <I>/D is observed at 22.3 and 25 keV. The trend of increasing diffracted intensity per unit absorbed dose as a function of energy was observed for all crystals: over 22 data series the mean increase in <I>/D between 12.4 and 25 keV is of 3.35 (figure 3). When using a detector with a silicon sensor, no such increase is observed with <I>/D approximately constant over the energy range used. Five additional data series were collected from two crystals over a higher X-ray energy range (22.3 - 27 keV) as the optimal energy for data collection using a CdTe based detector is expected to be at around 26 keV^31^ Over this limited energy range we observe no clear peak in the diffracted intensity per unit dose with <I>/D approximately constant between 22 and 26 keV, though a significant decrease, beyond experimental error, is observed at 27 keV as expected (figure 3 inset).

**Figure 3:**
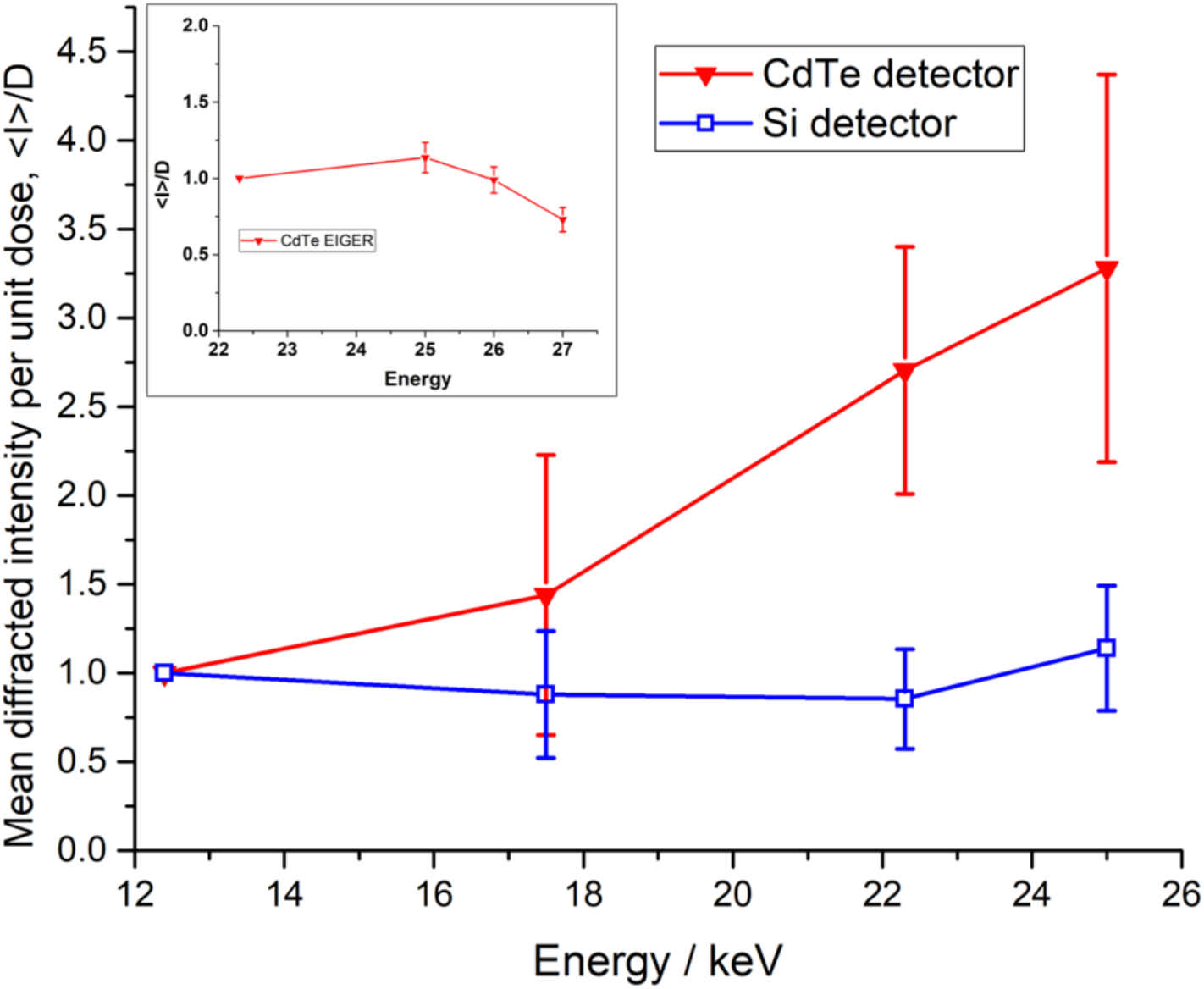
Increase in the diffracted intensity per unit absorbed dose, <I>/D, as a function of energy. Data shown are averaged from 29 data series (22 recorded using CdTe Eiger and 7 using the Si Pilatus) with the standard deviation at each energy shown as error bars. Variation in <I>/D over an energy range of 22.3-27 keV observed in additional 5 data series is shown in the inset. These data are normalised to 22.3 keV.

While an increase in diffraction efficiency as a function of energy is predicted by theory^13^, the size of the gain observed in these experiments is larger than predicted^31^ by a factor of ~2. There are other experimental factors which act to increase the gains realized for high energy data collection. Firstly, the beam used in these experiments has a Gaussian rather than top-hat profile. The theoretical increase of 1.6 in DE between 12.4 and 25 keV predicted by Dickerson and Garman^31^ assumes a top-hat beam, with gains of 3 to 4 only realized for beam and crystal sizes of less than 2 microns. When a Gaussian beam is used, photoelectrons will not be generated evenly within the illuminated volume, which is partly reflected in an increased theoretical diffraction efficiency of 1.9 between 12.4 and 25 keV when this is factored into the dose calculation. Some additional gain may therefore result as photoelectrons migrate out of the central high dose region, aided by the longer path lengths of photoelectrons at higher energies. Secondly, one has to consider that, despite best efforts, small errors in the measurement of the X-ray beam size and intensity and crystal size can easily result in under or over-estimation of the absorbed dose and hence I/D. We sought to minimise this as a source of error through the use of multiple crystals and data collection over multiple sessions with the flux and beamsize measured at each.

Some studies postulate that intensities at higher energies can be enhanced relative to lower energies due to a reduced background and lower absorption^38,39^. Here, we choose to keep the resolution at detector edge constant, moving the detector further away from the crystal at higher energies, since this is how crystallographers will best exploit the energy dependent gains for real-world data collection. This increase in crystal to detector distance has an influence on the data measured and an increase in both mean spot size and background intensity is observed with increasing energy. Spot size is heavily dependent on beam divergence, hence a longer sample-to-detector distance at higher incident beam energy will lead to a larger spot on the detector surface. With the CdTe Eiger detector an increase in measured background was seen at higher energies, consistent with elastic scattering from the non-crystalline component of the sample being the main factor as it is concentrated at smaller angles with increasing beam energy^26^. Our observations suggest that variation in spot size and background intensity are not significant influences on improvements in the metrics of diffraction data quality seen at higher energies.

In the above, photoelectron escape has not been considered when calculating absorbed doses. While this effect is negligible at 12.4 keV, at higher energies photoelectrons can escape from the illuminated volume reducing the effective deposited dose. At 25 keV the desposited dose is reduced by 20% for 20 μm crystals to D_PE_ (D and D_PE_ calculated using *RADDOSE-3D* and shown in table S2). Compared with three-fold increase in <I>/D shown in figure 3, <I>/D_PE_ shows a further increase as a function of energy, increasing by a factor of 4 between 12.4 and 25 keV (figure S4). Initial experiments with microcrystals of a volume smaller than the smallest available beamsize of 7 μm × 8 μm show that these effects amount can amount for a factor of 6 over this energy range (data not shown).

To experimentally confirm the predicted resolution dependence of the gains in diffracted intensity per unit dose, high resolution (1.25 Å) data series were also collected. The dimensions of the 9M Eiger and the currently achievable minimum crystal to detector distance at I24 preclude data collection to this resolution at 12.4 keV so these data series were normalized to 17.5 keV. Figure 4a shows the mean diffracting power per unit absorbed dose, <I>/D, in different resolution shells as a function of energy for data recorded using the CdTe Eiger. A clear resolution dependence is observed with <I>/D increasing by a factor of 2.7 for the lowest resolution shell compared to 4.2-fold over the range 1.35-1.25 Å. It follows that the high resolution limit of the diffraction data should also increase at higher energies. Figure 4b shows the energy dependent resolution cut-off for five crystals diffracting to between 2.1 and 1.65 Å at 12.4 keV. The resolution limit was determined using *dials.scale* applying a cut-off criterion of CC_1/2_ > 0.3. For all crystals, an energy dependent increase in the resolution cut-off is observed with a gain of between 0.2 and 0.3 Å. The resolution gain is also clearly visible in the electron density maps (figure S2). Reduced background and lower absorption at high X-ray energies also act to explain the gain in resolution.

**Figure 4:**
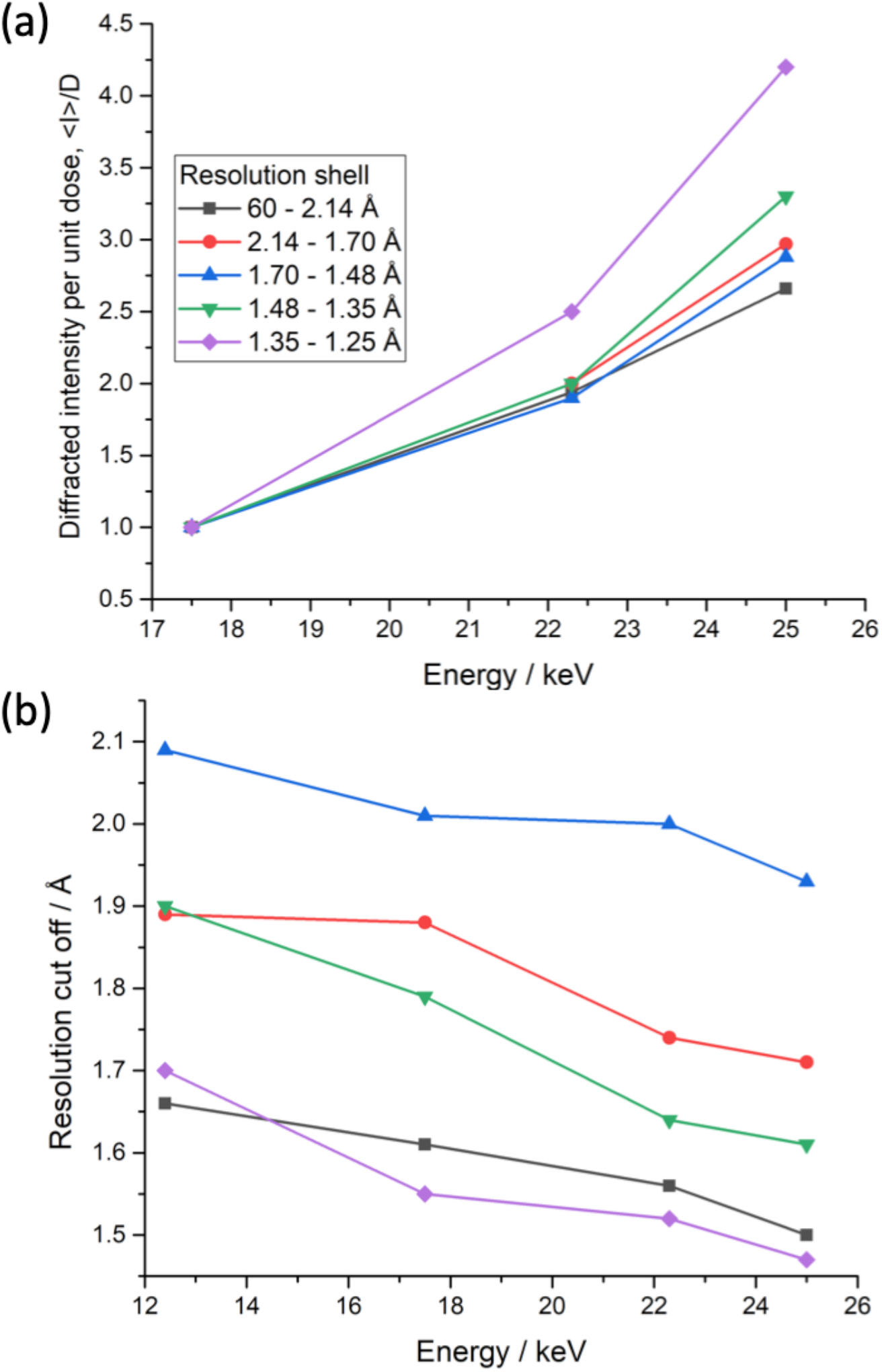
(a) Resolution dependence of the diffracted intensity per unit absorbed dose, <I>/D, and (b) energy dependent change in resolution cutoff as ascertained using a criterion of CC_1/2_ > 0.3.

With the advancement of synchrotron technology, from undulators providing high photon fluxes at high energies through to large area detectors able to efficiently record high-energy photons, it is now possible to efficiently and routinely collect data at energies which are optimal for macromolecular crystallography. Within this study, we show experimentally that a CdTe based detector enables the increased diffraction efficiency of crystals at high X-ray energies to be exploited due to its increased quantum efficiency in comparison to silicon-based detectors. The energy dependence of intensities follows Darwin’s law in both cases. The benefits of collecting at 25 keV include an increase in the resolution of data up to 0.3 Å that can be recorded for a given absorbed dose, compared to data collected at 12.4 keV. This increase in information content can for example enable the identification of water molecules in structural enzymology and make a critical difference in understanding structural function of proteins.

These results impact all macromolecular crystallography experiments from rotation to serial even when the crystal sizes used are relatively modest (~20 microns) and should be considered in the design of future MX beamlines and for X-ray data collection from all samples that yield crystals of limited size.

## Methods

### Sample preparation and sample mounting

Thermolysin crystals were grown in CrystalQuickX 96 well sitting drop plates (Greiner) using a Mosquito crystallization robot (STP Labtech) at 20°C. 50 mg/ml lyophilized thermolysin (Sigma) was dissolved in 0.05 M MES pH 6, 45 % DMSO, 50 mM NaCl and equilibrated against 1.2 M ammonium sulphate in a 1:1 ratio with a final drop size of 200 nl. 50% Ethylene glycol or 3 M ammonium sulphate were used as cryo-protectant. Crystals were mounted in loops and then cryo-cooled. Dimensions of the rod-shaped crystals were measured using the data acquisition GUI (GDA) and ranged from 20-40 μm in diameter with lengths of 120-310 μm.

### Beamline setup

All experiments were carried out at beamline I24 at Diamond Light Source. A new cryogenic permanent magnet undulator (CPMU) was installed shortly before the first experiments described here. Commissioning resulted in a varying X-ray flux between each experimental session. The X-ray flux was measured using a PD300-500 silicon PIN diode^40^ (Canberra), built by the Diamond Light source detector group and calibrated by the Physikalisch-Technische Bundesanstalt (PTB) up to energies of 60 keV, at the start of each session and is given in table S1. I24 features a two-stage focusing design with two pairs of Kirkpatrick-Baez mirrors, the first pair of mirrors features stripes with different coatings. For data collections below 20 keV a rhodium stripe was used, for experiments above 20 keV a platinum stripe were used. As the shape of mirrors was optimised at 12.4 keV, the beamsize increased when the Pt strip was used. The secondary pair of mirrors were not translated during the experiments as the Rh and Pt stripes overlap. Beam sizes at the sample position were determined by performing a knife-edge scan on a 200 μm thick gold wire and FWHM are given in table S1. Care was taken to measure beamline variables such as beamsize and the photon flux prior to each data collection session.

### Detector setup

Data were collected using an Eiger2 X 9M detector with a 750 μm thick CdTe sensor and a Pilatus3 X 6M with a 450 μm thick silicon sensor. The maximum frame rate of the Pilatus detector is 100 Hz (10 ms exposure times), while the Eiger detector can run at 230 Hz (4.3 ms). The detectors are mounted in an up and under configuration on a translation stage allowing data to be collected from each interchangeably. The sensor quantum efficiencies of each detector were interpolated from measured values provided by Dectris (Baden, Switzerland).

### Data collection

For each data series diffraction data were collected at 12.4, 17.5, 22.3 and 25 keV at a single position on a crystal. The order in which each energy was collected was varied between each data series. Doses were low and ranged between 300 and 800 kGy per dataset. At each position on a crystal repeated wedges of 100° of diffraction data were collected, with a single wedge at each energy. The crystal to detector distance was varied such that the inscribed circle on the detector face corresponded to the same resolution at all energies. When multiple data series were collected from a single crystal the crystal was translated by at least twice the beam FWHM between each series.

To enable direct comparison of the performance of the CdTe Eiger detector to that of the Si Pilatus detector, the exposure time was set to 10 ms in session A for the data collected at 22.3 keV or below. In subsequent sessions when the Pilatus was not used for comparison, an exposure time of 5 ms was used. In session A, at least one set of collections at the four energies was performed with both detectors from the same crystal. For the data at 25 keV, exposure times up to 83 ms at full transmission were required to compensate for the lower flux as well as the lower scattering efficiency. To be able to calculate the flux as accurately as possible, intensity values from a X-ray beam position monitor close to the sample position were recorded for all datasets. In session D, higher energies became accessible due to the ongoing commissioning of the CPMU, though exposures of 0.7 s per data frame were required at 27 keV. As commissioning continues it is anticipated that higher fluxes, and hence reduced exposure times, will become accessible with any time penalty associated with high energy data collection removed. To allow the energy dependence of <I>/D at high resolution to be probed, exposure times were increased and a shorter crystal to detector distance used during this session.

### Data processing

Data were processed with *DIALS* v.3.0.4 integration package^41^. To generate statistics with consistent parameters across data sets, *xia2*^42^ was used. Raw intensities were used from the data processing integration step to avoid complications introduced by scaling routines or inadvertent ‘scaling out’ of energy dependent differences.

Crystals typically diffracted to 1.4-2.0 Å, taking into account diffraction in the corner of the detector and applying the CC_1/2_ > 0.33 criterion. However, a resolution cut-off of 2.0 Å was applied to all data when quantifying energy dependent changes in the mean diffracting power. This was defined by the maximum resolution of the inscribed circle at the minimum detector distance at 12.4 keV.

The average diffraction weighted dose, referred to as dose here, was calculated for each dataset using *RADDOSE-3D*^43^. Dose calculations assumed that the long axis of the crystal is oriented vertically towards the X-ray beam. The <I>/D value is defined as to the unscaled mean intensity given by *xia2* from the lowest resolution shell up to 2.0 Å divided by the deposited dose estimated for the individual data set.

## Acknowledgements

We would like to thank Halina Mikolajek and Sam Horrell for producing the thermolysin crystals. Andrew Foster, James O’Hea, Scott Williams and Adam Taylor are thanked for setting up the CdTe Eiger detector on I24. Furthermore, we thank Colin Nave for his very helpful comments on the manuscript and James Holton for enlightening discussions on Darwin’s equation.

## Author contributions

S.L.S.S. planned the experiment with contributions from R.L.O. and D.A. The data collection was performed by all authors. Most of the data analysis was done by S.L.S.S. with support from R.L.O and D.A‥ The manuscript was written by S.L.S.S., D.A. and R.L.O.

## Supplementary material

**Table S1:** Overview of all diffraction data collected. A data series refers to a set of four datasets collected at four different energies from a single position on a crystal. Doses were calculated using *RADDOSE-3D* taking photoelectron escape into account.

| Session & number of data series | Energy / keV | Incident flux / $\times 10^{11}$ ph s$^{-1}$ | Beamsize / $\mu\text{m}^2$ | Crystal diameter / $\mu\text{m}$ | Total exposure time per dataset / s | Total deposited dose per dataset / kGy |
| --- | --- | --- | --- | --- | --- | --- |
| **A**<br>8 CdTe<br>7 Si | 12.4 | 26.40 | $7.9 \times 8.9$ | 25 | 10 | 280 |
|  | 17.5 | 8.48 |  | 25 | 10 | 440 |
| | 22.3 | 4.09 | $9.7 \times 10.2$ | 25 | 10 | 350 |
|  | 25.0 | 0.255 |  | 25 | 83 | 400 |
| **B**<br>8 CdTe | 12.4 | 52.80 | $7.0 \times 8.1$ | 25 | 5 | 390 |
|  | 17.5 | 17.00 |  | 25 | 5 | 590 |
| | 22.3 | 4.80 | $9.0 \times 10.6$ | 25 | 5 | 510 |
|  | 25.0 | 0.32 |  | 25 | 70 | 500 |
| **C**<br>6 CdTe | 12.4 | 46.8 | $7.9 \times 9.1$ | 20 | 5 | 400 |
|  | 17.5 | 18.0 |  | 20 | 5 | 570 |
| | 22.3 | 4.52 | $9.0 \times 10.7$ | 20 | 5 | 590 |
|  | 25.0 | 0.32 |  | 20 | 70 | 530 |
| **D**<br>5 CdTe | 22.3 | 5.06 | $9.0 \times 10.7$ | 25 | 5 | 580 |
|  | 25 | 0.31 |  | 25 | 70 | 510 |
|  | 26 | 0.26 |  | 25 | 90 | 470 |
|  | 27 | 0.04 |  | 25 | 700 | 500 |
| **E**<br>5 CdTe | 17.5 | 18.0 | $7.9 \times 9.1$ | 30 | 8 | 840 |
| | 22.3 | 4.52 | $9.0 \times 10.7$ | 30 | 8 | 840 |
|  | 25 | 0.32 |  | 30 | 110 | 700 |

**Table S2:** Data collection parameters and integration statistics obtained for the two data series shown in figure 2, S1 and S2 and S3. Note that<I> and <I/σ(I)> reported are for unscaled data allowing direct comparison between independent datasets. CC_1/2_ and R_meas_ reported are for scaled data as conventionally reported as indicators of dataset quality. All statistics are given in the form (overall, low resolution, high resolution). Statistics given are for over the entire resolution range with values in brackets are for the outermost resolution shell (2.0-2.03 Å). The resolution cut was set to 2Å, as the inscribed circle at 12.4 keV is limited to this resolution by the detector area. The resolution given in the table was calculated based on the CC_1/2_ >0.3 criterion and takes reflections in the corners of the detector into account. The dose refers to the diffraction weighted dose obtained using RADDOSE3D as discussed in the text. The data collection sequence for both crystals was 12.4 keV, 17.5 keV, 22.3 keV, 25 keV

| Data series | 1 | 1 | 1 | 1 | 2 | 2 | 2 | 2 |
| --- | --- | --- | --- | --- | --- | --- | --- | --- |
| Energy | 12.4 | 17.5 | 22.3 | 25.0 | 12.4 | 17.5 | 22.3 | 25.0 |
| Detector | CdTe Eiger | CdTe Eiger | CdTe Eiger | CdTe Eiger | Si Pilatus | Si Pilatus | Si Pilatus | Si Pilatus |
| Detector QE | 0.912 | 0.933 | 0.940 | 0.945 | 0.835 | 0.475 | 0.264 | 0.193 |
| Resolution range / Å | 60-2.0, 60-5.4, 2.03 -2.0 |  |  |  |  |  |  |  |
| N obs | 239882, 12598, 12117 | 239717,12433, 12160 | 239887,12401, 12341 | 243415, 12520, 12379 | 326214, 16409, 15841 | 328963,16457, 16139 | 238551, 11683,12174 | 240791, 11715, 12298 |
| <I> | 6.5, 11.5, 1.6 | 11.1, 20.0, 2.3 | 10.5, 18.8, 2.4 | 16.2, 29.0, 3.7 | 5.0, 8.9, 1.3 | 3.9, 7.0, 1.0 | 4.1, 7.3, 1.0 | 6.3, 11.4, 1.6 |
| <I/σ(I)> | 9.7, 24.1, 2.9 | 10.5, 26.9, 2.5 | 8.9, 22.6, 2.4 | 11.0, 27.1, 3.1 | 8.3, 20.5, 2.5 | 6.3, 15.9, 1.8 | 5.0, 13.0, 1.3 | 6.2, 15.7, 1.8 |
| R<sub>meas</sub> | 0.191, 0.115, 0.570 | 0.171, 0.101, 0.542 | 0.196, 0.110, 0.604 | 0.116, 0.068, 0.352 | 0.244, 0.114, 0.910 | 0.303, 0.131, 1.167 | 0.316, 0.147, 0.945 | 0.249, 0.119, 0.733 |
| R<sub>pim</sub> | 0.135, 0.081, 0.403 | 0.121, 0.072, 0.383 | 0.139, 0.078, 0.427 | 0.116, 0.068, 0.352 | 0.074, 0.037, 0.275 | 0.092, 0.041, 0.350 | 0.096, 0.047, 0.282 | 0.079, 0.040, 0.229 |
| CC<sub>1/2</sub> | 0.958, 0.965, 0.640 | 0.969, 0.978, 0.686 | 0.961, 0.976, 0.622 | 0.977, 0.979, 0.714 | 0.992, 0.992, 0.814 | 0.989, 0.994, 0.710 | 0.986, 0.991, 0.791 | 0.990, 0.993, 0.861 |
| Incident flux ph/s | 2.22 × 10<sup>10</sup> | 7.56× 10<sup>10</sup> | 1.37 × 10<sup>11</sup> | 2.54 × 10<sup>10</sup> | 2.88 × 10<sup>10</sup> | 7.88 × 10<sup>10</sup> | 1.35 × 10<sup>11</sup> | 2.76 × 10<sup>10</sup> |
| total exposure / s | 5 | 5 | 5 | 83 | 5 | 5 | 5 | 83 |
| Dose / kGy | 290 | 480 | 410 | 490 | 380 | 500 | 400 | 540 |
| Deposited dose /kGy | 280 | 440 | 350 | 400 | 370 | 460 | 350 | 430 |

**Figure. S1:**
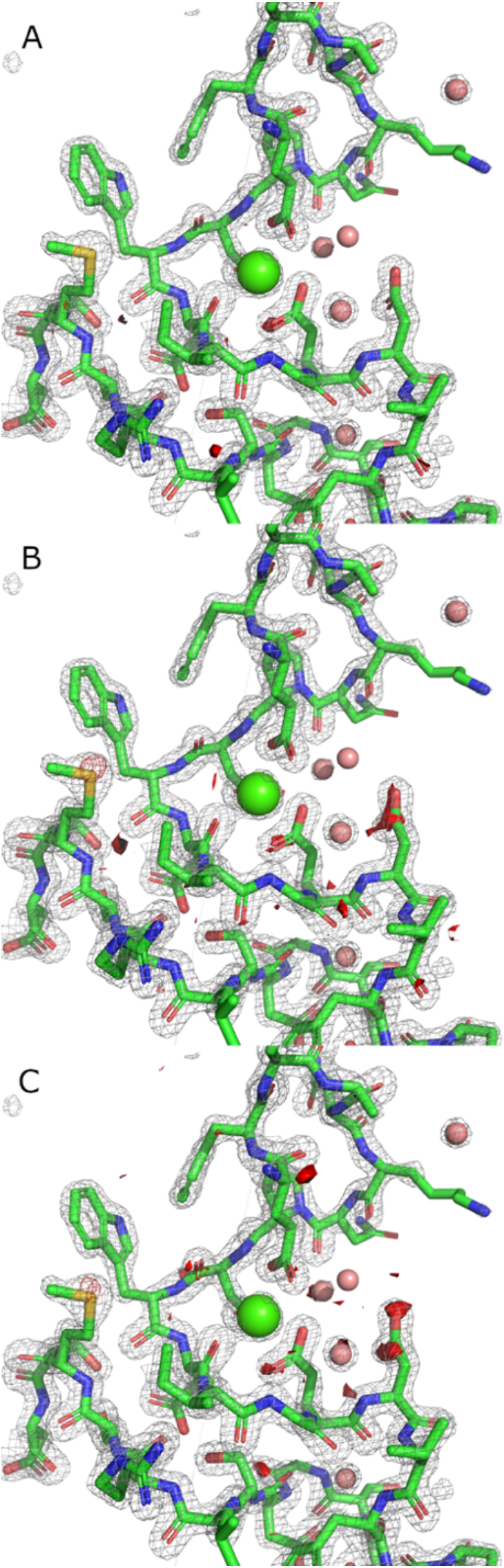
Exemplar electron density obtained from sequential datasets forming a data series. The grey mesh represents the 2Fo-Fc map and the stick model the refined thermolysin structure from the first dataset in the series (17500 eV). Data were recorded using the CdTe Eiger detector and processed using phenix.refine (Liebschner *et al*. (2019)). These data were collected in the order: 17500eV, 22300 eV, 25000 eV, 12400 eV. In A, B, and C Fo-Fo difference maps have been superimposed onto the refined data, shown in opaque surface representation and contoured to 4 sigma. A shows 12400eV minus 17500eV, B shows 12400eV minus 22300eV, C shows 12400eV minus 25000eV. Maps produced with phenix.fobs_minus_fobs_map. Very slight density loss is visible on acidic residues ASP190, ASP191 and the sulphur atom of MET205 in the third and fourth datasets collected.

**Figure. S2:**
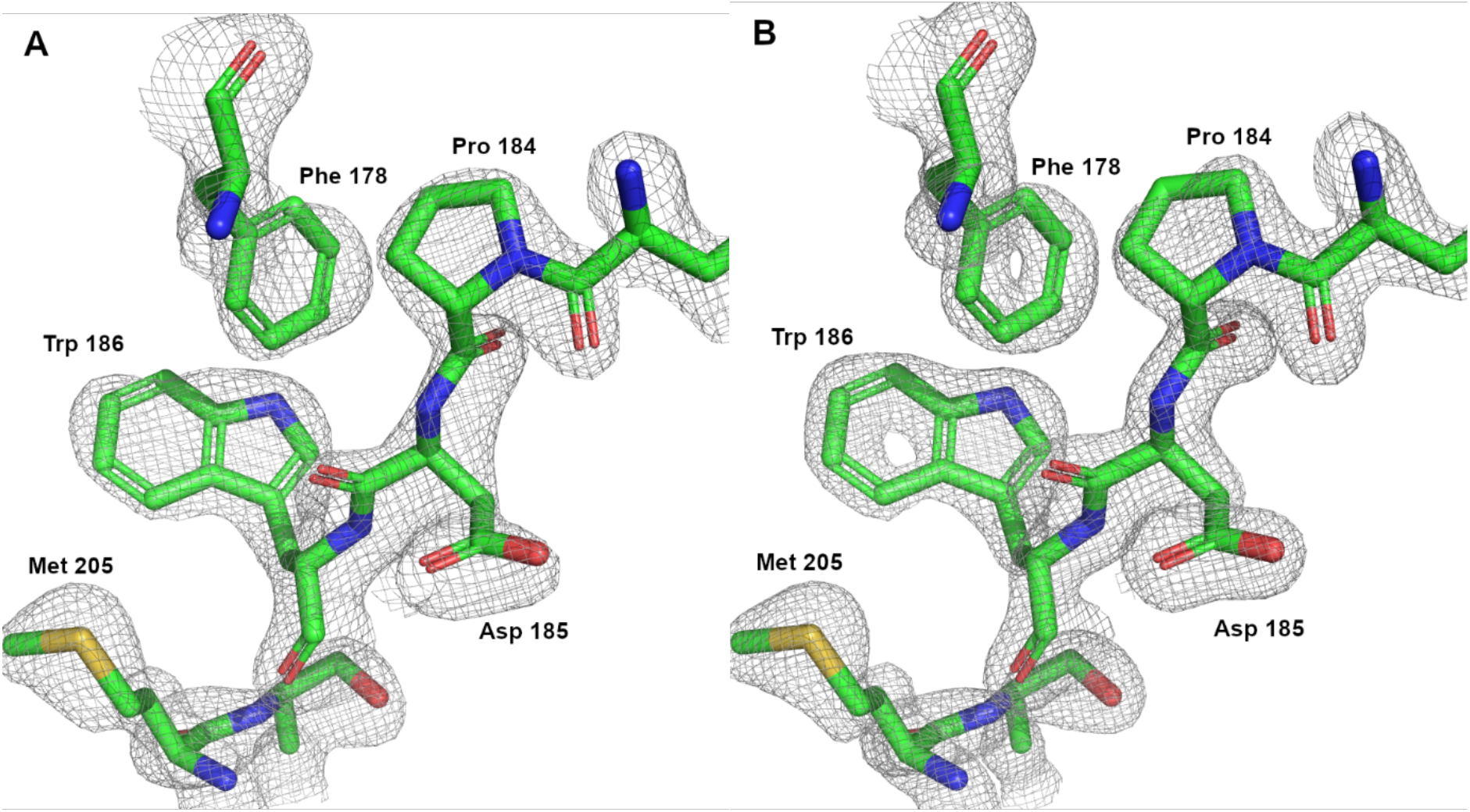
Composite omit maps illustrating differences between data recorded using the CdTe detector at different energies but the same total diffracted intensity. (A) Example volume of electron density obtained from 100 degrees of thermolysin data collected at 12.4 keV, diffracting to 1.90 Å using a CC_1/2_>0.3 threshold. (B) electron density of the same volume obtained by collecting from the same position of the crystal at 25 keV and diffracting to 1.61 Å using the same critereon. The increased resolution is apparent in the electron density of rings and clarity of other features. Maps contoured to 1.5 σ in both cases. Models were refined and maps obtained using Phenix; R/R_free_ 0.210/0.245 in the case of (A) and R/R_free_ 0.202/0.229 in the case of (B). Figure created using Pymol (The PyMOL Molecular Graphics System, Version 2.0 Schrödinger, LLC).

**Figure S3:**
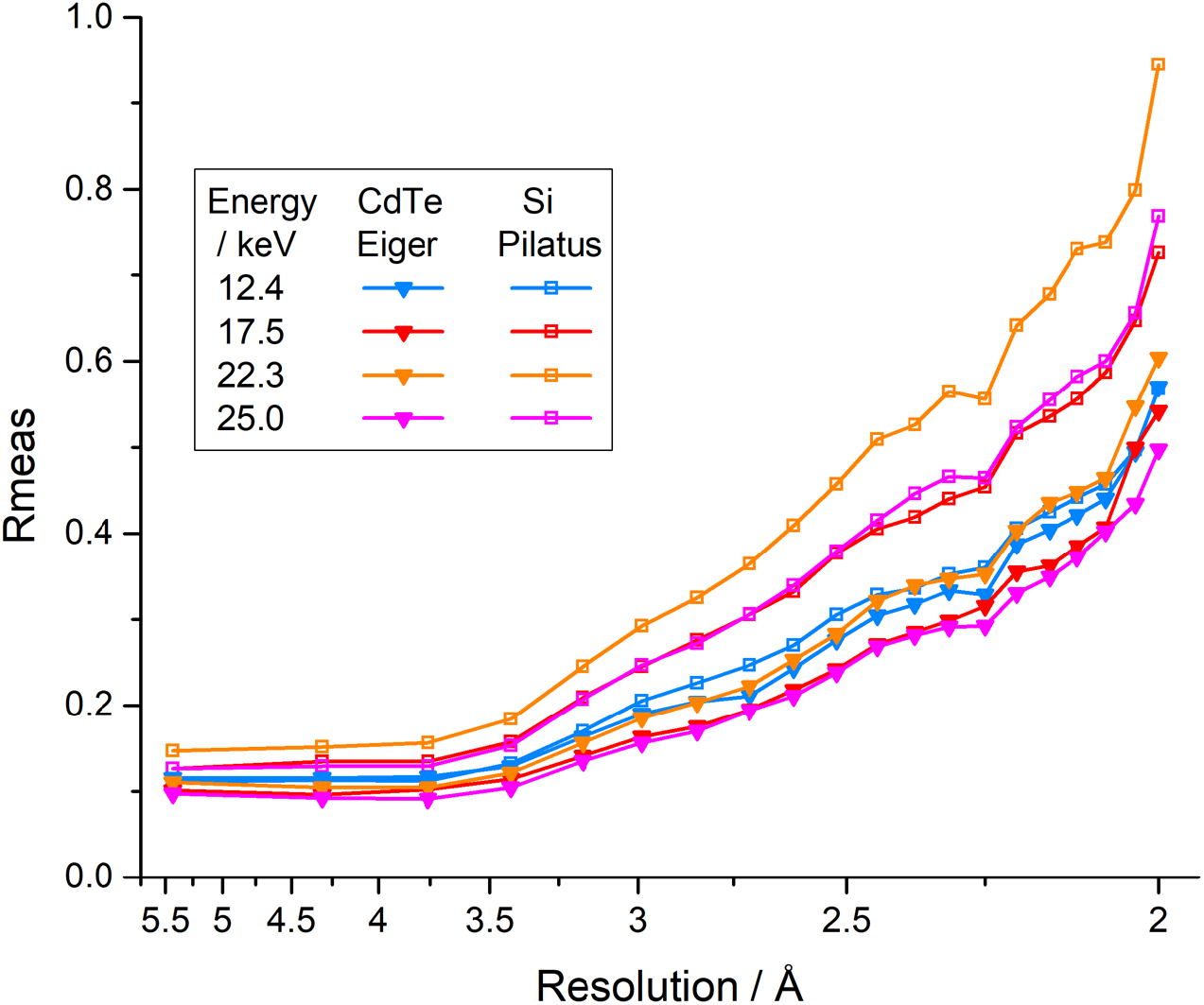
Redundancy independent merging R value R_meas_ for the data series summarized in figure 2 and table S2.

**Figure S4:**
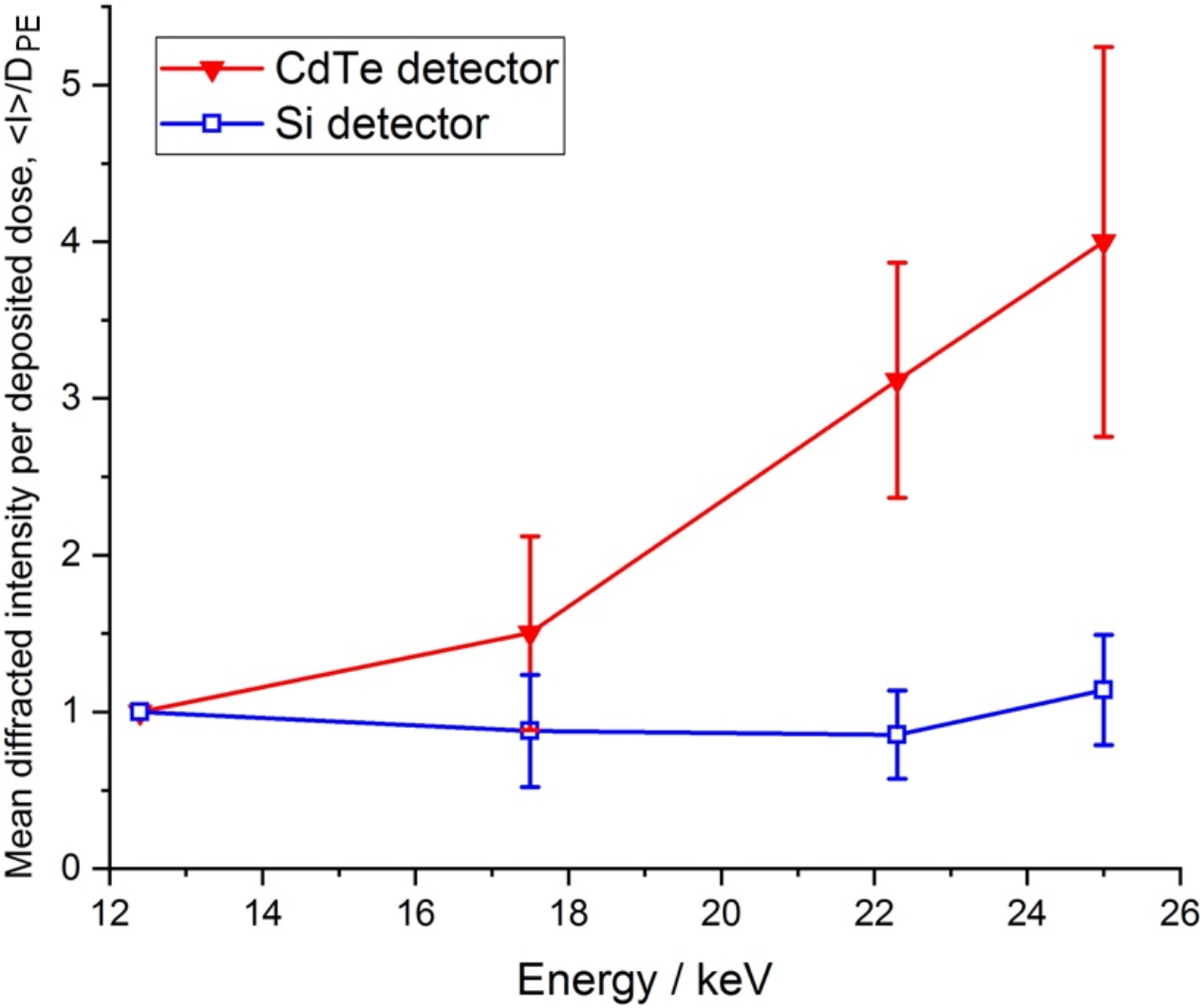
Increase in the diffracted intensity per unit deposited dose, <I>/D_PE_, as a function of energy. Data shown are averaged from 29 data series (22 recorded using CdTe Eiger and 7 using the Si Pilatus) with the standard deviation at each energy shown as error bars.

## References appearing in supplementary material only

D. Liebschner, P. V. Afonine, M. L. Baker, G. Bunkóczi, V. B. Chen, T. I. Croll, B. Hintze, L.-W. Hung, S. Jain, A. J. McCoy, N. W. Moriarty, R. D. Oeffner, B. K. Poon, M. G. Prisant, R. J. Read, J. S. Richardson, D. C. Richardson, M. D. Sammito, O. V. Sobolev, D. H. Stockwell, T. C. Terwilliger, A. G. Urzhumtsev, L. L. Videau, C. J. Williams, and P. D. Adams Acta Cryst. (2019). D75, 861-877

